# Growth of Mesenchymal Stem Cells at the Surface of Silicone, Mineral and Plant-Based Oils

**DOI:** 10.1101/2022.08.17.504254

**Authors:** Lihui Peng, Julien E. Gautrot

## Abstract

Bioemulsions are attractive platforms for the expansion of adherent cells in bioreactors. Their design relies on the self-assembly of protein nanosheets at liquid-liquid interfaces, displaying strong interfacial mechanical properties and promoting integrin-mediated cell adhesion. However, most systems developed to date have focused on fluorinated oils, which are unlikely to be accepted for direct implantation of resulting cell products for regenerative medicine, and protein nanosheets self-assembly at other interfaces has not been investigated. In this report, the composition of aliphatic pro-surfactants palmitoyl chloride and sebacoyl chloride, on the assembly kinetics of poly(L-lysine) at silicone oil interfaces and characterisation of ultimate interfacial shear mechanics and viscoelasticity is presented. The impact of the resulting nanosheets on the adhesion of mesenchymal stem cells (MSCs) is investigated via immunostaining and fluorescence microscopy, demonstrating the engagement of the classic focal adhesion-actin cytoskeleton machinery. The ability of MSCs to proliferate at the corresponding interfaces is quantified. In addition, expansion of MSCs at other non-fluorinated oil interfaces, based on mineral and plant-based oils is investigated. Finally, the proof-of-concept of such non-fluorinated oil systems for the formulation of bioemulsions supporting stem cell adhesion and expansion is demonstrated.

## 1. Introduction

The culture, scale up of manufacturing and processing of adherent cells requires innovative solutions to enable translation of emerging cell-based technologies in the fields of regenerative medicine and cultivated meat [1–2]. Although a range of platforms have been designed and proposed to answer these challenges, from hollow fibre reactors to plastic and hydrogel microcarriers for stirred tank reactors [3–6], these remain associated with complex cell processing steps, as well as high costs and risks of contamination with microplastics.

In this context, bioemulsions are attractive alternatives to solid microcarriers for the culture of adherent cells. Oil phases that could be applied are inexpensive and cell products can readily be separated and processed for further applications [7–9]. Recently, the proof-of-concept of long term expansion of mesenchymal stem cells at the surface of bioemulsions was demonstrated [7]. MSCs were cultured from passage 3 to passage 10 on bioemulsions, solid microcarriers and 2D tissue culture plastic in parallel, followed by phenotypic characterisation at multiple time points (cell morphology, flow cytometry, qPCR, tri-lineage differentiation assays). The phenotype of MSCs was found to be undistinguishable upon expansion in these three culture systems, at any given round of expansion tested, and instead was found to be sensitive to the passage number and prior rounds of population doubling.

Although the culture of adherent cells may seem surprising, owing to the wealth of reports linking a broad range of cell phenotypes to substrate mechanics [10–12], it was demonstrated that this phenomenon is enabled by the assembly of mechanically strong protein nanosheets at the liquid-liquid interface [13–14]. Indeed, corresponding interfaces display high interfacial shear moduli and elasticity, enabling resistance to cell-mediated contractile forces. Keese and Giaever had identified that surfactant molecules, combined to poly(L-lysine) enabled the culture of fibroblasts at the surface of oil droplets [15] and such phenomenon was recently assigned the formation of protein nanosheets [13]. More systematic studies of the combined impact of surfactant and protein chemistry on the co-assembly of nanosheets at fluorinated oil interfaces identified that covalent coupling of pro-surfactant molecules to associated proteins enabled the formation of quasi-2D (6-10 nm) elastic networks and the storage of strain energy upon cell-mediated contractile forces [16]. The impact of the molecular structure, including molecular weight and charge density, of proteins assembling to such interfaces on the toughness and viscoelasticity of resulting networks was also recently reported [17]. However, although fluorinated oil systems used in these studies have played important roles in discovery, analysis and diagnostics platforms in biotechnologies, such as microdroplet microfluidics [18–19], they remain unlikely solutions for implantation and cultured meat products. Other oil alternatives are therefore important to identify.

The present work explores the replacement of fluorinated oils and pro-surfactants with silicone and aliphatic oils, including mineral and rapeseed oils used routinely in cosmetics, consumer healthcare and the food industry. The formation of poly(L-lysine) nanosheets assembled at corresponding oil interfaces in the presence of the pro-surfactants hexadecanoyl chloride and sebacoyl chloride (hydrolysing into biocompatible palmitic acid and sebacic acid [20–21], respectively) is explored first via interfacial rheology, enabling the quantification of interfacial shear moduli, the kinetics of nanosheet formation and the investigation of resulting viscoelastic profiles of interfaces. The adhesion and expansion of mesenchymal stem cells cultured at resulting interfaces is then investigated. Finally, the stability of emulsions formed with corresponding protein nanosheets is investigated and the feasibility of MSC culture at corresponding non-fluorinated bioemulsions demonstrated.

## 2. Materials and Methods

### Materials

PBS, sodium hydroxide, poly(L-lysine) (PLL) hydrobromide (molecular weight 30 70 kDa), toluene (anhydrous, 99.8%), DMSO, triethylamine, triethoxy(octyl) silane, sebacoyl chloride (SBC), hexadecanoyl chloride (HDC), bovien serum albumin, gelatin (from cold water fish skin), fibronectin (from human plasma), paraformaldehyde, Triton X 100, tetramethyl rhodamine isothiocyanate phalloidin, 4,6 diamidino-2 phenylindole and monoclonal anti vinculin antibody from mouse were purchased from Sigma Aldrich. Live/Dead kit was from Life Technologies. Glass coverslip (25 × 60 mm) and ethanol were from VWR. PDMS (trimethylsiloxy terminated, viscosity 10 cSt) was from ABCR. Alexa Fluor 488 goat anti rabbit IgG (H+ highly cross adsorbed secondary antibody, Alexa Fluor 488 goat anti mouse IgG (H+ highly cross adsorbed secondary anti body, Hoechst 3334333422 (1(1 mg/mL)mg/mL) were purchased from ThermoFisher Scientific. Sticky-slide well was from Thistle Scientific. Rapeseed oil was from Sainsbury’s. Blandol^®^ White Mineral Oil was purchased from Sonneborn..

### Interfacial rheology

Interfacial rheology was carried out on a hybrid rheometer (DHR-3) from TA Instruments fitted with a double wall ring (DWR) geometry and a Delrin trough with a circular channel. The double wall ring used for this geometry has a radius of 34.5 mm and the thickness of the Platinum–Iridium wire is 1 mm. The diamond-shaped cross-section of the geometry’s ring provides the capability to pin directly onto the interface between two liquids and measure the interface properties without sub-phase correction. 19 mL of PBS (pH adjusted to 10.5) was placed in the Delrin trough and the ring was lowered, ensuring contact with the surface, via an axial force procedure. The measuring position was set 500 μm lower than the contact point of the ring with the aqueous phase surface. Thereafter, 15 mL of the oil phase containing the pro-surfactant mixture (hexadecanoyl chloride or sebacoyl chloride at final concentrations of 10 or 100 mg/mL, as specified in corresponding figures) was carefully syringed on top of the aqueous phase.

Time sweeps were performed at a constant frequency of 0.1 Hz and a temperature of 25 °C, with a displacement of 1.0 10^-3^ rad, to follow the formation of the protein layers at the interface. The concentration of poly(L-lysine) (PLL) used for all rheology experiments were 100 μg/mL (with respect to aqueous phase volume). Before and after each time sweep, frequency sweeps (with a constant displacement of 10^-3^ rad) were conducted to examine the frequency-dependant characteristics of the interface. Amplitude sweeps (with constant frequencies of 0.1 Hz) were carried out to ensure that the selected displacement was within the linear viscoelastic region. Considering the low moduli initially measured for pristine liquid-liquid interfaces (in the absence of protein and/or surfactant), viscous drag from both phases were not corrected. We note that although the interfacial shear moduli observed at liquid-liquid interfaces in the absence of protein or surfactant are expected to be considerably lower than those measured in our assay [59, 60], due to lack of viscous drag correction, they should not completely vanish and interfacial shear viscosity should be expected to persist, even with liquid-liquid interfaces that display very limited roughness (at the molecular scale).

Stress relaxation experiments were performed at 1% strain for 120 s. Stress relaxation data from the 10^th^ second onwards was plotted as a function of time and fitted with a double exponential decay function, according to the following equation:

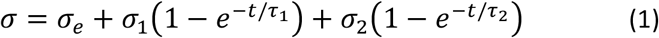

In this equation, *σ* is the measured residual stress, *σ_e_* is the elastic stress and *σ_1_* and *σ_2_* are viscous relaxation components. *τ_1_* and *τ_2_* are relaxation time constants. The degree of stress retention (*σ_r_*) is calculated as:

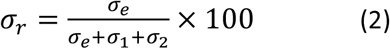

### Generation of pinned droplets

Glass samples were placed into vials containing toluene (1 mL), 30 μL trichlorooctyl silane and 30 μL triethylamine and incubated at room temperature (protocols were carried out in jars with sufficient volumes of toline/trichlorooctyle silane/triethylamine solutions, in order to produce multiple slides in parallel). After 24 h incubation, the glass slides were washed with ethanol and dried in air. The resulting hydrophobic glass slides were cut into 1×1 cm samples and placed into a 24-well plate. After sterilization with 70% ethanol, samples were washed with PBS and filled with 1 mL PBS (with pH adjusted as indicated). 100 μL pinned droplets of oil (silicone, mineral or rapeseed oil) containing the desired co-surfactant at 10 or 100 μg/mL concentrations were deposited on top of the submerged coated glass slide. 1 mL of 200 μg/mL PLL solution was pipetted into each well (final concentration of 100 μg/mL) and left to incubate for 1 h. Samples were washed by successive dilution/aspiration with PBS (pH 7.4, 6 times). Fibronectin adsorption was carried out by adding 20 μL of a fibronectin solution (1 mg/mL) into each well (final concentration: 10 μg/mL), followed by incubation at room temperature for 1 h. Finally, samples were washed by dilution/aspiration with PBS (pH 7.4) four times and then with growth medium twice.

### Preparation of glass-mounted wells for higher resolution imaging

Hydrophobized thin glass slides (25×60 mm) were attached to Sticky-Slide 8 Well (An 8 well bottomless μ-Slide with a self-adhesive underside to which substrates can be mounted, Ibidi). After sterilization with 70% ethanol, wells were washed with PBS and filled with 600 μL PBS (with pH adjusted as indicated). Then 10 μL pinned droplets of the desired oil supplemented with the co-surfactant (HDC and/or SBC; concentrations specified in corresponding figures) were deposited on top of the submerged coated glass substrates. 300 μL PBS was removed by micropipette aspiration. 300 μL of 200 μg/mL PLL solution was pipetted into each well (final concentration of 100 μg/mL) and left to incubate for 1 h. Each well was washed by successive dilution/aspiration with PBS (pH 7.4, 6 times). Fibronectin adsorption was carried out by adding 6 μL of a fibronectin solution (1 mg/mL) into each well (final concentration: 10 μg/mL), followed by incubation at room temperature for 1 h. Finally, each well was washed by dilution/aspiration with PBS (pH 7.4) four times and then with growth medium twice.

### Formation of bioemulsions

Oils were pre-mixed with 100 μg/mL of the corresponding surfactants (HDC or HDC/SBC mixtures) and added into a glass vial that had been plasma oxidized for 10 minutes. A PLL solution (200 μg/mL in pH 10.5 PBS; volume double that of the oil phase) was then pipetted into the vial, prior to vortexing for a few seconds. After an hour incubation, the bottom phase below the emulsion level was gently aspirated using a syringe with a long needle and PBS was slowly injected to dilute unreacted proteins. The process was repeated four times to dilute the excess PLL and normalise the pH.

### Mesenchymal stem cells (MSCs) culture and seeding

Bone marrow derived human mesenchymal stem cells (PromoCell) were cultured on T75 flasks in MSC growth medium (PromoCell). MSCs were harvested with 4 mL accutase-solution (PromoCell), resuspended, then centrifuged. 5,000 cells per well (resuspended in medium) were seeded on flat interfaces (per well in 24 well plates) and cultured in an incubator (37°C and 5% CO2). Half of the medium was replaced with fresh medium every two days. For passaging, 300,000 cells were seeded in a T75 flask.

### Hoechst staining and cell counting

Cell proliferation on flat interfaces was assessed via Hoechst staining. Half of the medium was replaced by pre-warmed PBS containing 2 μL Hoechst (1 mg/mL Thermofisher Scientific). After 30 min incubation, cells were imaged using a Leica DMI4000 fluorescence or or Leica DMi8 epifluorescence microscope. Cell counting was carried out by thresholding and watershedding nuclear images in ImageJ.

### Cytocompatibility assay

Cell viability was assessed via Live/Dead staining. Half of the medium was replaced with pre warmed PBS containing 1 μL/mL calcein, 4 μL/mL ethidium from Live/Dead kit (Life Technologies) and 2 μL/mL Hoechst. After 40 min incubation, cells were imaged using a Leica DMI4000B epifluorescence microscope. Cell counting was carried out by thresholding and watershedding nuclear images via ImageJ.

### Immunostaining

For immunostaining and imaging at higher resolution, cells were cultured on pinned droplet generated in a sticky-slide 8 well plates (Ibidi), prepared as described above. After 24 or 48 h incubation, each well was diluted with PBS six times before samples were fixed with 8 % paraformaldehyde for 10 min and diluted with PBS six times before permeabilization with 0.4% Triton X-100 for 5 min at room temperature. Samples were blocked for 1 h (blocking buffer: PBS containing 10 vol% foetal bovine serum and 0.5 vol% gelatine), combining with tetramethyl rhodamine isothiocyanate phalloidin (1:500, Sigma-Aldrich). Samples were subsequently incubated with primary antibodies (anti-vinculin mouse monoclonal, Sigma-Aldrich, 1:200 in blocking buffer) for 1 h at room temperature, diluted with PBS six times, then incubated with Alexa Fluor 488-conjugated secondary antibodies (goat anti-mouse, 1:500 in blocking buffer) and DAPI (1:500) for 1 h at room temperature. The samples were washed six times by dilution with deionised water and imaged shortly after.

### Immuno-fluorescence microscopy and data analysis

Fluorescence microscopy images were acquired with a Leica DMI4000B fluorescence microscopy (CTR4000 lamp; 63 × 1.25 NA, oil lens; 10 × 0.3 NA lens; 2.5 x 0.07 NA lens; DFC300FX camera) and a Leica DMi8 epifluorescence microscope (HC PL FLUOTAR 10x/0.32 PH1; HC PL FLUOTAR 63X/1.30 Oil PH3; LEICA DFC9000 GT sCMOS camera). Confocal microscopy images were acquired with a Leica TCS SP2 confocal microscope (X-Cite 120 LED lamp; 63 x 1.40-0.60 NA, oil lens; 10 x 0.3 NA lens; DFC420C CCD camera) and a Zeiss Super resolution LSM710 ELYRA PS.1 (EC Plan-Neofluar10x/0.3 M27; EC Plan-Neofluar20x/0.5 M27; sCMOS camera). Cell densities were determined after thresholding and watershedding nuclei images in ImageJ. In the case of cell aggregates, cells were counted manually. To determine adhesion cell areas, images were analysed by outlining the contour of the cell cytoskeleton (phalloidin stained) and areas were measured in ImageJ. To determine cell spreading areas, images were analyzed by thresholding and watershedding cytoskeleton images (phalloidin staining). For confocal imaging, stacks of 16 sections were scanned, with an image averaging of 2 and a line averaging of 4. 3D reconstruction and volume rendering of the stacks were performed via Imaris x64.

### Statistical analysis

Statistical analysis was carried out using Origin 2019 through one-way ANOVA with Tukey test for posthoc analysis. Significance was determined by * P < 0.05, ** P < 0.01, *** P < 0.001 and n.s., non-significant. A full summary of statistical analysis is provided below as a separate supporting file.

## 3. Results and Discussion

In order to promote cell adhesion and expansion at the surface of non-fluorinated liquids, we proposed to adapt the formulation of protein nanosheets previously investigated at fluorinated interfaces [16, 22]. In these systems, the cationic protein poly(L-lysine) (PLL) was introduced in the aqueous phase (PBS) and allowed to co-assemble with the pro-surfactant pentafluorobenzoyl chloride (PFBC) introduced in the oil phase. We referred to this latter molecule as pro-surfactant as neither PFBC, nor PLL, on their own display sufficient tensio-active properties to stabilise emulsions. However, when combined, these two molecules can couple and form a layer of protein that is physically crosslinked at the interface between the two liquids. Covalent coupling of pro-surfactant to proteins such as PLL, albumin or lysozyme was found to be essential to confer sufficient elasticity to the interface to store energy upon deformation and resist cell-mediated contraction [16].

Therefore, we translated such design to the formation of PLL nanosheets at non-fluorinated oil-water interfaces, combining PLL with aliphatic acyl chlorides. Two pro-surfactants were selected (Figure 1): hexadecanoyl chloride (HDC), displaying a long aliphatic tail and hydrolysing into the food-compatible and biocompatible palmitic acid [23–24], and sebacoyl chloride (SBC) [20–21], enabling direct intermolecular covalent crosslinking within nanosheets and also hydrolysing into cytocompatible residues (sebacic acids, anhydrides and esters enter the composition of drug release systems) [16, 20]. As in the case of nanosheet assembly at fluorinated interfaces, the pH of the aqueous phase was adjusted to 10.5, corresponding to the pKa of PLL, to enable sufficient coupling of acyl chloride residues [17, 22].

**Figure 1.**
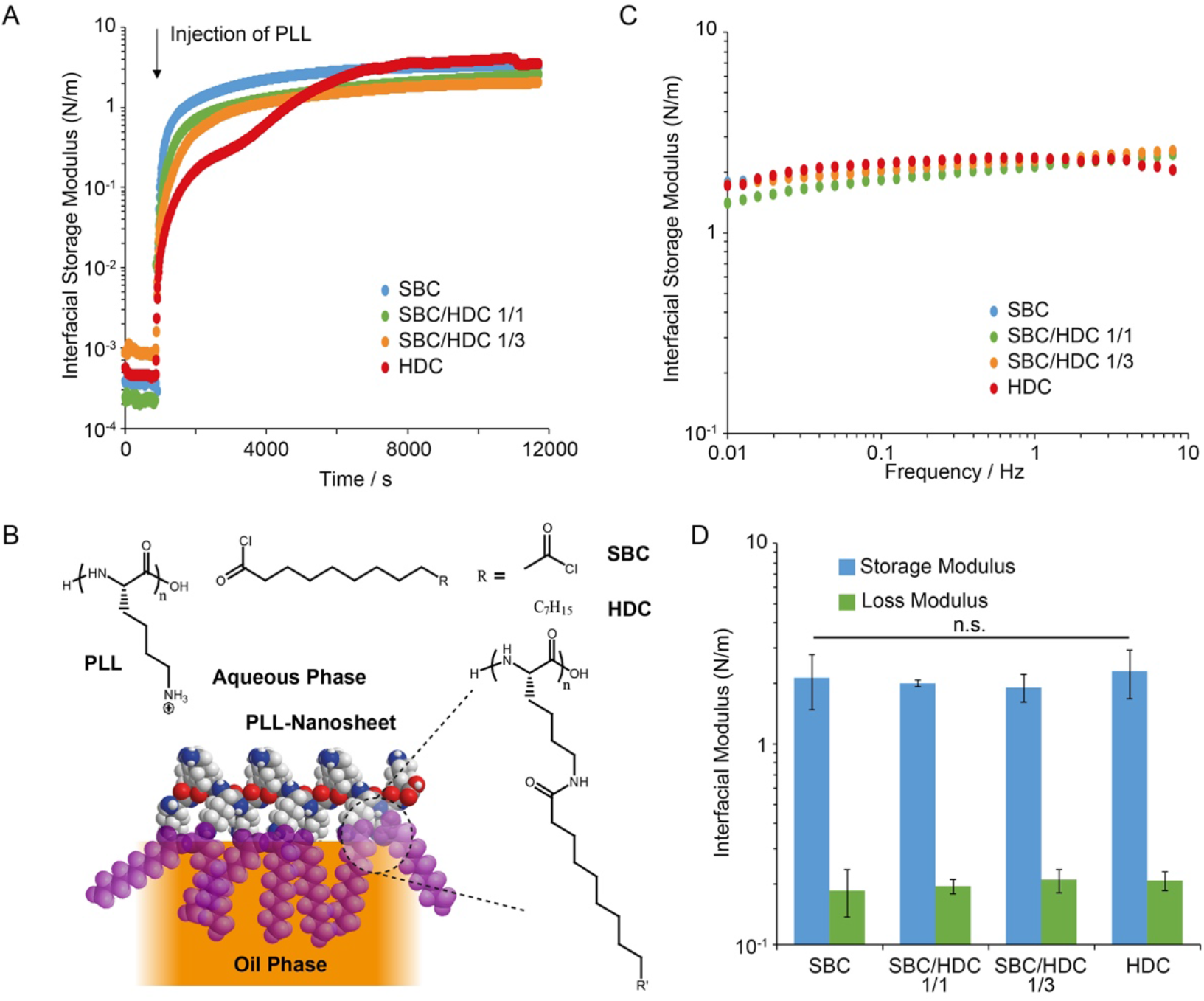
Pro-surfactant assisted PLL assembly forms a mechanically strong viscoelastic nanosheet at liquid PDMS interfaces. A) Representative interfacial time sweep rheology profiles recorded at a frequency of 0.1 Hz and strain of 1% ([PLL] = 100 μg/mL; [pro-surfactant] = 10 μg/mL; weight ratios are reported for mixed surfactants). B) Chemical structure and proposed idealised architecture of protein nanosheets. R’ correspond to the heptadecyl chains, in the case of HDC, and amide crosslinks or unreacted carboxylic acids in the case of SBC. C) Representative frequency sweep profiles for different pro-surfactant combinations, recorded at a strain of 1 %. D) Corresponding interfacial storage and loss moduli measured at 0.1 Hz (error bars are s.e.m.; n=3).

The assembly of PLL nanosheets at silicone oil interfaces was first examined, using interfacial rheology (Figure 1A). Time sweeps demonstrated the rapid increase in interfacial shear moduli associated with the self-assembly of PLL-pro-surfactant nanosheets at silicone-water interfaces. Upon introduction of PLL, both the storage (Figure 1A) and loss moduli (Supplementary Figure S1) rapidly increased, and diverged as the storage moduli rapidly dominated. Moduli gradually levelled to a plateau, with storage moduli in the range of 1-2 N/m. Hence these data confirm the rapid formation of protein networks at interfaces, associated with > 3 orders of magnitude increase in interfacial shear moduli. Such phenomenon is comparable, in terms of kinetics and magnitude, to the self-assembly of PLL nanosheets at fluorinated oil interfaces [16, 22].

Although the pro-surfactant formulation had no significant impact on the ultimate modulus of PLL nanosheets (Figure 1D), it impacted the kinetics of the evolution of interfacial storage moduli (Figure 1A). As HDC content increased, the storage moduli increased more slowly as a function of time. In addition, in the case of nanosheets assembled in the presence of HDC alone, a two-step process was observed. After a first plateau seemed to be reached near 2000 s post injection, around 3000 s a second increase was observed. This could reflect the formation of bilayers or simply the more gradual shift from a soft physically crosslinked nanosheet to a more structured bilayer (see proposed schematic representation of such bilayer in Figure 1B). Indeed, evidence for such bilayer was obtained in similar systems (with hexadecane as the oil) for PLL-HDC nanosheets, from neutron reflectometry experiments [16]. The longer alkyl chains associated with HDC, compared to PFBC and SBC (able to generate strong physical and covalent crosslinks respectively), may be associated with the initial softer character of the interfaces observed and the requirement for relatively high functionalisation levels to achieve comparable interfacial moduli. However, it should be noted that PLL surface densities and HDC/SBC functionalisation levels were not quantified in the present specific system.

Further analysis of the frequency sweeps revealed comparable viscoelastic profiles (Figure 1C), in agreement with the establishment of elastic networks at corresponding interfaces. Little frequency dependency was observed over the range probed (0.01-10 Hz). Storage moduli were consistently well above loss moduli (approximately one order of magnitude), further supporting the elastic nature of nanosheets formed. Overall, all pro-surfactant formulations resulted in nanosheets with comparable moduli and viscoelastic profiles (Figures 1C-D). Interfacial elasticity is an important property that was recently proposed to play an essential role in enabling cell adhesion at liquid-liquid interfaces [16]. Therefore establishing a detailed understanding of the viscoelastic profile of PLL nanosheets at non-fluorinated oil interfaces is particularly important.

To further investigate interfacial viscoelasticity, stress relaxation experiments were carried out at interfaces. In such experiments, the du Noüy ring allowed the establishment of a controlled strain on corresponding nanosheets, and the relaxation of interfacial stress was monitored as a function of time. The relaxation profiles were then fit with a double exponential decay function, in order to extract stress retention σ_r_ and relaxation rate constants [16]. Relaxation profiles recorded for PLL nanosheets formed in the presence of different pro-surfactant formulations were comparable, almost overlapping (Figure 2A). In turn, the stress retentions and relaxation constants measured remained within narrow ranges (σ_r_ between 74 and 80% and τ_1_ between 73 and 91 s; Figure 2B and Supplementary Figure S2). These results confirm the high level of elasticity associated with PLL nanosheets formed at silicone-water interfaces, for all pro-surfactant combinations tested. Therefore, PLL nanosheets formed with aliphatic pro-surfactants mediating physical or covalent crosslinks displayed combinations of high interfacial shear moduli and elasticity that are expected to support cell adhesion, spreading and proliferation at liquid interfaces.

**Figure 2.**
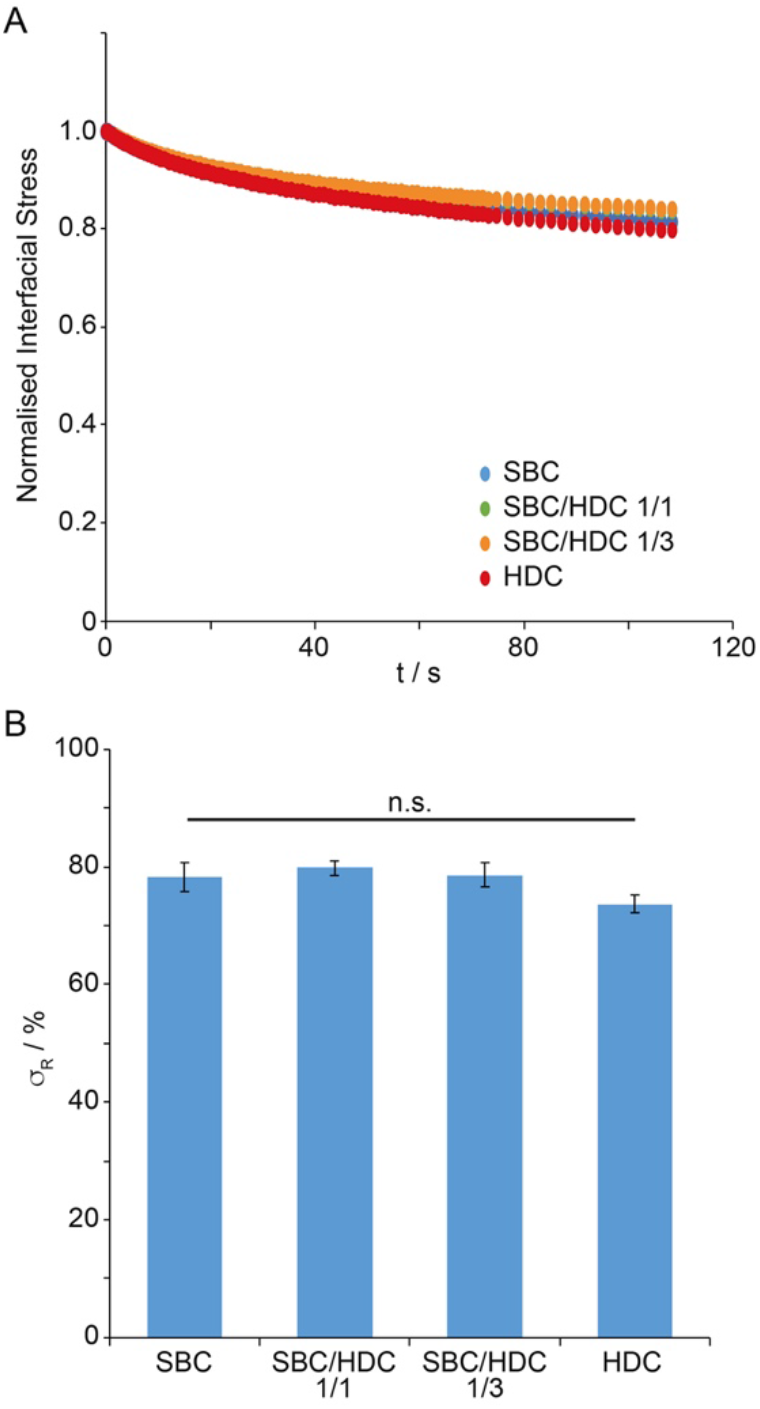
Viscoelasticity of PLL nanosheets formed at liquid PDMS interfaces. A) Representative stress relaxation profiles for different pro-surfactant combinations, recorded after 1% strains ([PLL] = 100 μg/mL; [pro-surfactant] = 10 μg/mL; weight ratios are reported for mixed surfactants). B) Stress retentions (σ_R_) extracted from corresponding double exponential fits (error bars are s.e.m.; n=3).

Indeed, a broader screen of different protein and pro-surfactant formulations had indicated that the elasticity of protein nanosheets assembled at liquid-liquid interfaces was essential to sustain cell-mediated spreading and expansion, in the context of primary keratinocytes, mesenchymal stem cells and the epidermal cell line HaCaT [16]. Therefore the viscoelastic properties of PLL nanosheets assembled at silicone interfaces in the presence of HDC and SBC pro-surfactants suggest that these interfaces should enable the adhesion, spreading and expansion of MSCs at corresponding non-fluorinated oil-water interfaces. It is proposed that such combination of high interfacial shear modulus and capacity to store strain energy upon cell-mediate deformation will resist cell mediated contractility and sustain cell spreading.

MSCs were seeded at PLL nanosheet-stabilised interfaces formed in the presence of SBC/HDC mixtures, further coated with fibronectin, and allowed to spread for 24 to 48 h. The spreading of resulting cells was evaluated by immunostaining and microscopy (Figure 3). At both time points, MSCs spreading on PLL nanosheets at silicone oil interfaces could be found to display spread morphologies with total surface areas comparable to those of cells spreading on glass coverslip coated with PLL and fibronectin (2000-2700 μm^2^). In all cases, cells displayed lamellipodia associated with fan-shaped morphologies and numerous focal adhesions, identified from vinculin stainings. MSCs also displayed well developed actin stress fibres connecting to focal adhesions. Therefore, our data indicate that MSCs spread at PLL nanosheet-stabilised silicone oil interfaces via the classic integrin-actomyosin machinery, in good agreement with observations of primary keratinocytes and MSCs spreading on fluorinated oil interfaces stabilised by PLL-PFBC assemblies [13, 16].

**Figure 3.**
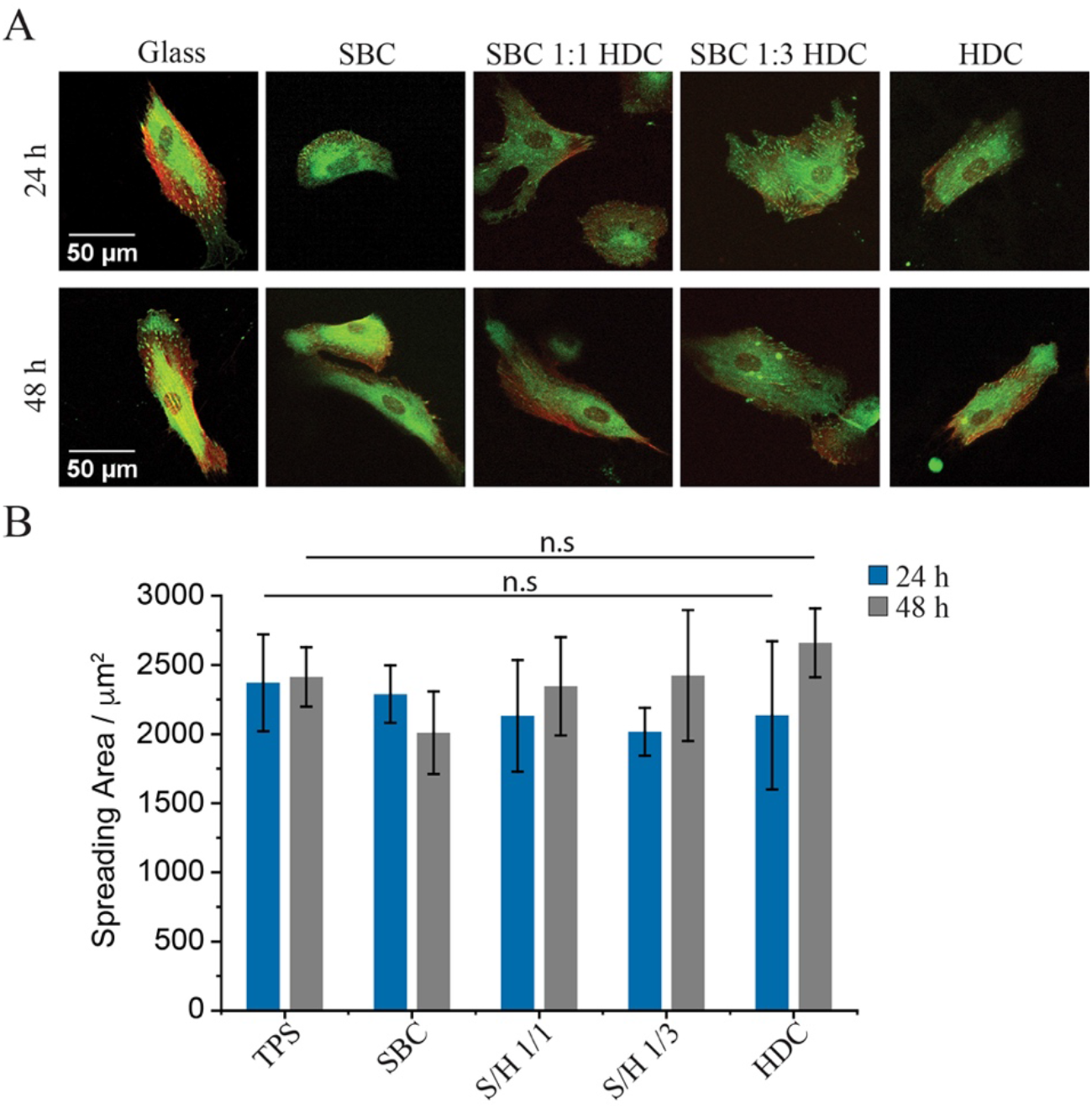
Impact of pro-surfactant composition on MSC spreading at liquid silicone interfaces. MSCs were cultured at the surface of PLL (100 μg/mL) nanosheets formed in the presence of pro-surfactant mixtures at 10 μg/mL, followed by functionalisation with fibronectin (20 μg/mL), at liquid silicone interfaces. A) Confocal microscope images of MSCs cultured at the interfaces for 24 hours (top) and 48 hours (bottom), green, vinculin; red, actin. B) Spreading area quantified from corresponding images and conditions. Error bars are s.e.m,; n ⩾ 3.

The proliferation of MSCs at silicone oil interfaces was next examined. The cytocompatibility of these liquid-liquid culture systems was investigated first at two different pro-surfactant concentrations (Figure 4). At concentrations of 10 μg/mL, cell viabilities remained comparable on all liquid interfaces tested, as well as the tissue culture polystyrene (TPS) control, both at days 1 and 3 (Figure 4A). Cell densities were also comparable on all interfaces, at early and late time points (days 3 and 5), indicating that initial cell adhesion and proliferation remained comparable on PLL nanosheet-stabilised interfaces and rigid tissue culture plastic substrates. At higher pro-surfactant concentrations (100 μg/mL), although cell densities and viabilities were found to be lower on pure sebacoyl chloride/PLL and hexadecanoyl chloride/PLL nanosheets, mixed pro-surfactant/PLL assemblies displayed comparable cytocompatibility and capacity to sustain expansion to the TPS controls (Figure 4B). These results are in good agreement with previous observations that the pro-surfactant pentafluorobenzoyl chloride induced some cytotoxicity at concentrations above 10 μg/mL [13], although this effect was more pronounced in the case of PFBC. Therefore, although changes in local pH associated with the hydrolysis of residual acyl chloride is likely to play a role in such phenomenon, the inherent toxicity of perfluoroaromatics, compared to palmitic acid and sebacic residues, may also contribute to such effects.

**Figure 4.**
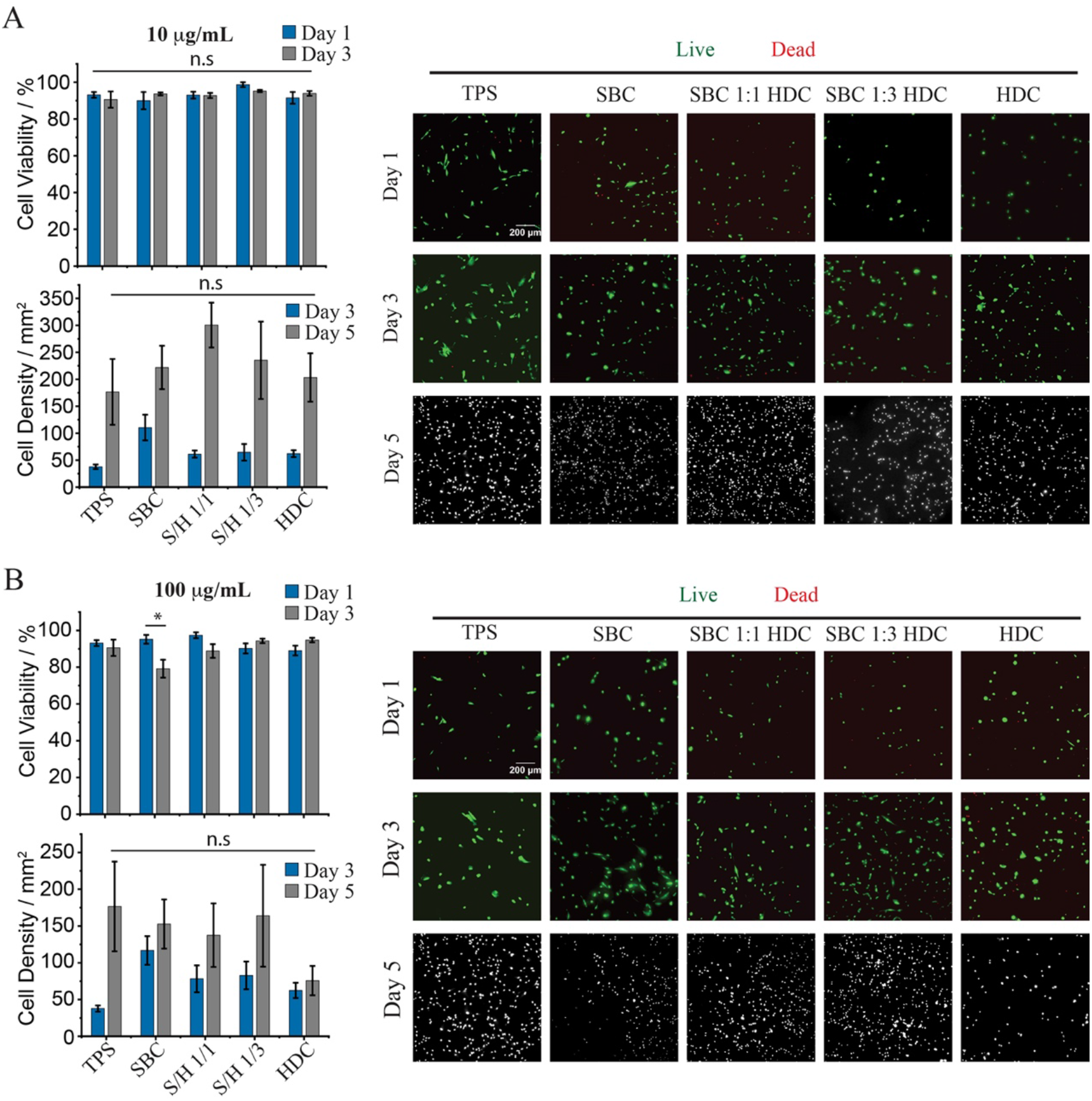
Impact of pro-surfactant composition on MSC proliferation at liquid silicone interfaces. MSCs were cultured at the surface of PLL (100 μg/mL) nanosheets formed in the presence of pro-surfactant mixtures at 10 (A) and 100 μg/mL (B), followed by functionalisation with fibronectin (20 μg/mL), at liquid silicone interfaces. Summary of viabilities and cell densities (left, top and bottom, respectively) and corresponding epifluorescence microscopy images (right). Error bars are s.e.m,; n ⩾ 3.

To further demonstrate the translatability of aliphatic pro-surfactant/PLL nanosheets for the culture of MSCs at the surface of a broader range of oils, the ability of MSCs to expand on two aliphatic oil substrates was next examined (Figure 5). In particular, we selected a mineral oil used in the formulation of consumer health products and rapeseed oil, typically used as cooking oil, as aliphatic oil case studies, owing to their inherent cytocompatibility and potential for translation in cell manufacturing platforms. Early stage (day 3) cell densities were comparable on both mineral and rapeseed oils to those achieved on TPS. After 6 days of culture, cell densities on PLL nanosheets stabilised with SBC/HDC 1/1 were slightly below those measured on TPS, although these differences were not statistically significant in the experimental conditions tested. Note that cell densities on the control used (TPS) differ in Figures 5A and b as the passage numbers used in corresponding experiments differed. Overall, our data indicate that PLL nanosheets stabilised by mixtures of palmitoyl chloride and sebacoyl chloride enable MSC adhesion and expansion on a broad range of non-fluorinated oil interfaces.

**Figure 5.**
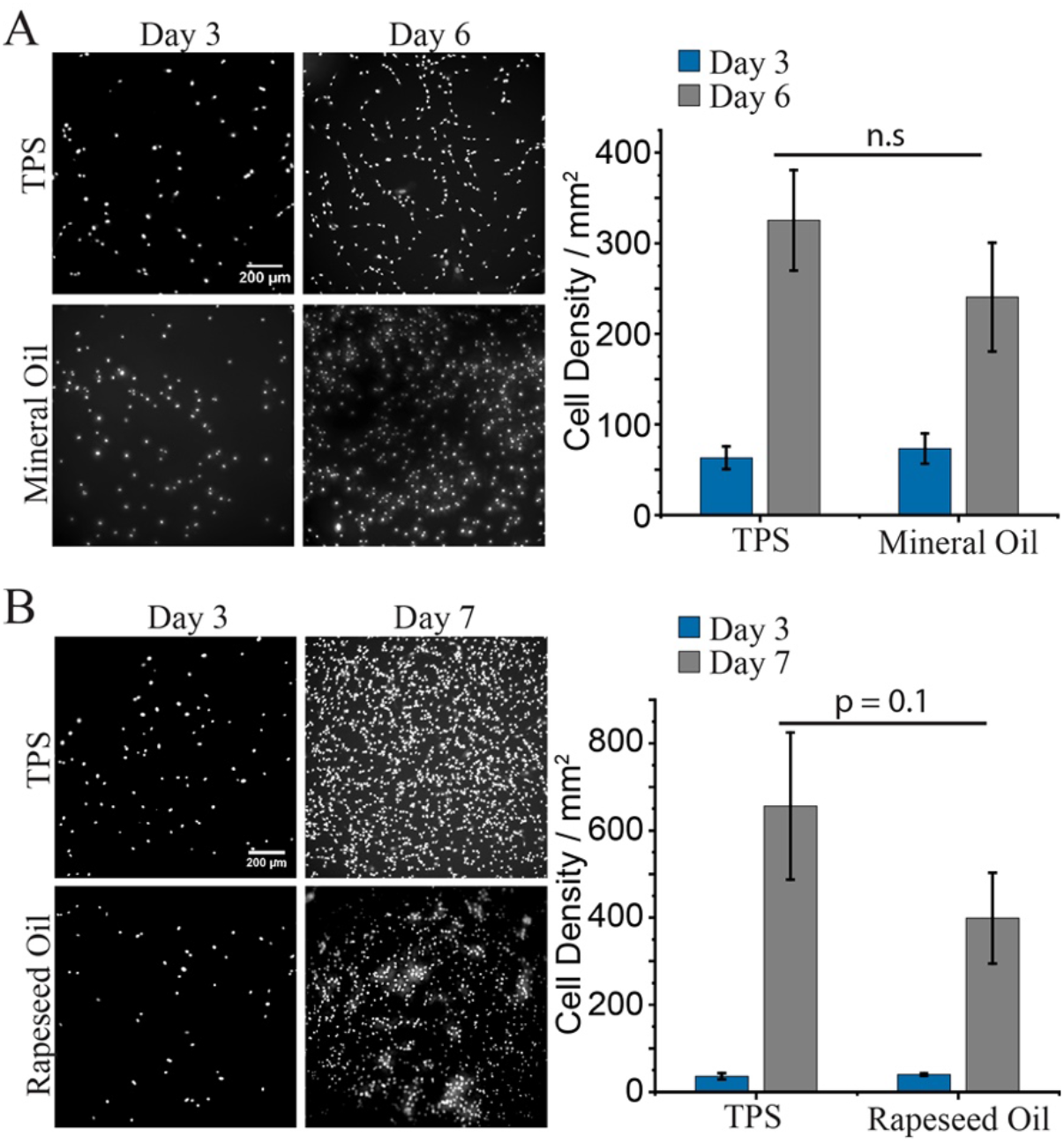
Proliferation of MSCs on aliphatic oils (mineral and rapeseed oils). MSCs were cultured at the surface of PLL (100 μg/mL) nanosheets formed in the presence of HDC at 10 μg/mL, followed by functionalisation with fibronectin (20 μg/mL), at mineral and rapeseed oil interfaces. Epifluorescence microscopy images (left) of MSC nuclei on (A) mineral oil and (B) rapeseed oil interfaces, and corresponding cell densities (right). Error bars are s.e.m,; n ⩾ 3

Finally, the ability to expand MSCs at the surface of non-fluorinated oil emulsions was explored. The stability of emulsions stabilised by different pro-surfactant combinations was first examined (Figure 6A and Supplementary Figure S3). Whereas PLL-based emulsions stabilised by HDC-rich pro-surfactants were stable just after formation and washing (removal of excess PLL), and after 24 h, those formed with SBC-rich nanosheets rapidly destabilised, in particular those based purely on SBC. Hence, the greater conformational freedom associated with longer aliphatic chains of palmitoyl residues, and their reduced degree of tethering with PLL chains enabled the improved stabilisation of non-fluorinated oil emulsions.

Finally, MSCs were seeded at the surface of mineral oil and silicone emulsions stabilised with PLL/fibronectin nanosheets, and allowed to expand for 7 days. The resulting cultures were then stained with calcein to identify viable cells. MSCs could be seen covering the surface of corresponding microdroplets, with many cells clearly spreading. Therefore, these results are in good agreement with the cell expansion observed at corresponding 2D protein nanosheet stabilised liquid-liquid interfaces. Together, these results indicate that aliphatic oil emulsions could constitute attractive adherent cell culture platforms for the expansion of stem cells in bioreactors for cell manufacturing.

**Figure 6.**
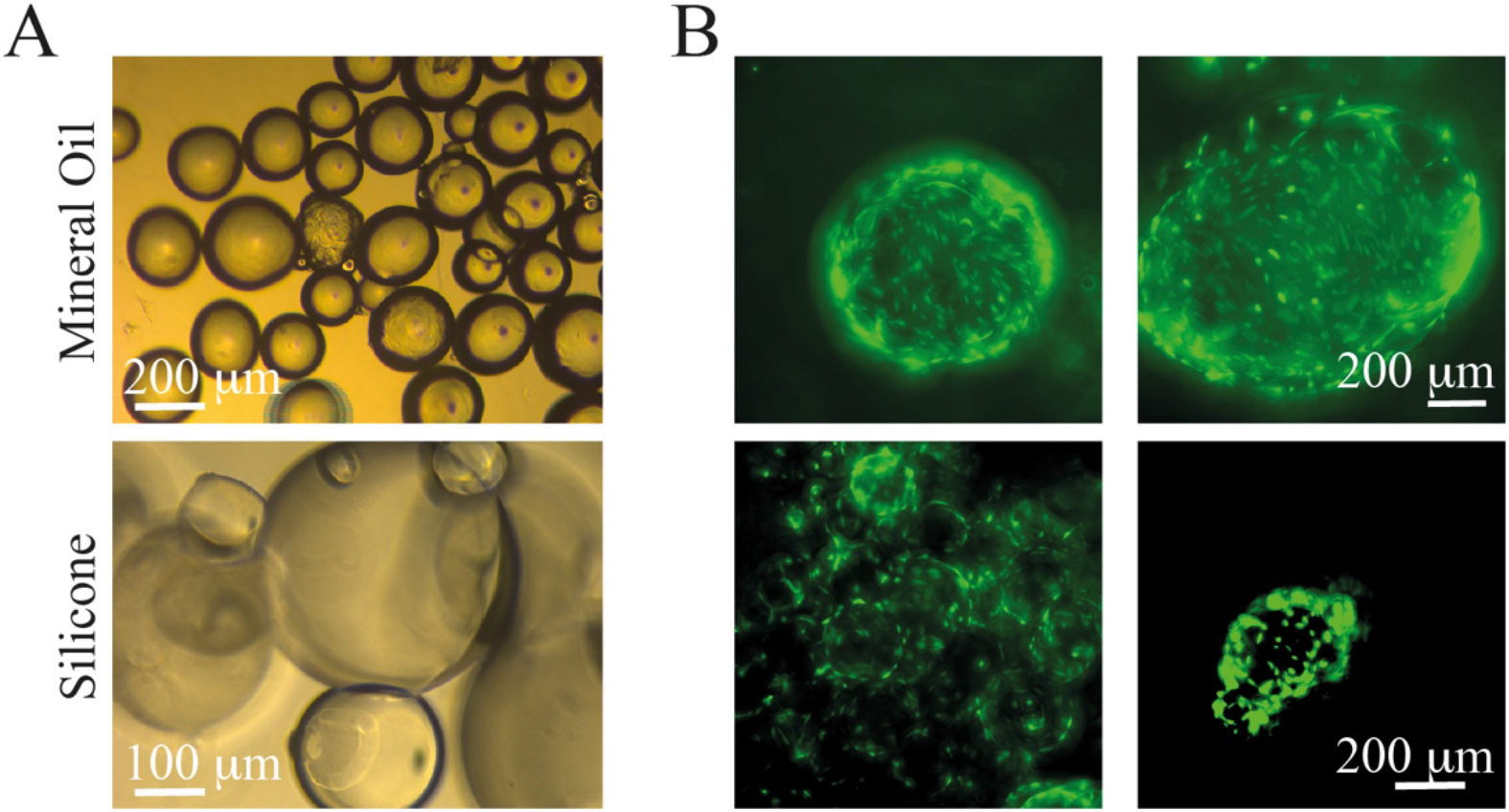
MSC cultured on non-fluorinated oil-based emulsions. A) Bright field microscope images of mineral (top) and PDMS emulsions (bottom) incubated for 3 days. B) Epifluorescence microscopy images of MSCs cultured on corresponding emulsions for 7 days (calcein staining). Emulsions were prepared using the corresponding oils supplemented with 100 μg/mL SBC/HDC 1/1, functionalised with PLL/FN nanosheets (PLL, 100 μg/mL; fibronectin, 20 μg/mL).

## 4. Conclusions

Cells sense the biochemical and physical properties of their environment. Mechanical engagement of integrins with surrounding protein networks, strengthening of focal adhesions and assembly of the actin cytoskeleton requires a balance of forces, supported by macromolecular networks, sensed at the nanoscale [25]. The dimensions of focal adhesions and actin stress fibres, in the range of a few tens of nanometres, and their mechanical properties, inherent of self-assembled soft matter systems, are comparable with those of other protein nanostructures, including protein nanosheets at liquid-liquid interfaces. Therefore, strong PLL networks displaying interfacial moduli in the range of 1-2 N/m and thicknesses in the range of 10-20 nm, even in the case of non-fluorinated oil interfaces and pro-surfactants, are well within the range of dimensions and mechanical properties typically associated with the integrin-talin-vinculin-acto-myosin machinery. Nanoscale sensing of mechanical properties therefore further appears as a key element for the design of new generations of biomaterials. The results presented demonstrate that such concepts and design principles are not restricted to fluorinated oil interfaces, and can be more broadly implemented to other interfaces, including silicone and aliphatic oils. The combined role of pro-surfactants and PLL chemistry on the self-assembly of nanosheets displaying strong mechanical properties and high elasticity remains central to the ability to support cell adhesions and expansion. Specifically, MSCs seeded on PLL nanosheet-stabilised silicone interfaces developed focal adhesions and assembled mature actin cytoskeletons. MSCs were found to proliferate to comparable levels on such interfaces to cells cultured on tissue culture polystyrene. In turn, aliphatic oil interfaces, reinforced using equivalent assemblies, enabled the expansion of MSCs too and supported their culture on emulsions. This proof-of-concept, although calling for systematic characterisation and quantification of cell phenotype at non-fluorinated bioemulsion interfaces, demonstrates the feasibility of stem cell cultures at the surface of mineral and plant-based oils that are relevant to consumer healthcare and cultured meat industry.

## Supporting Information

Supporting Information is available from the author.

## Acknowledgements

Funding for this work from the European Research Council (ProLiCell, 772462; ProBioFac, 966740), the China Scholarship Council (studentship, 201708060335) and the Leverhulme Trust Foundation for financial support (RPG-2017-229, Grant 69241) is gratefully acknowledged.

