## Supplementary Information for "Growth of Mesenchymal Stem Cells at the Surface of Silicone, Mineral and Plant-Based Oils"

**Lihui Peng[1,2] and Julien E. Gautrot[1,2*]**

*[1] Institute of Bioengineering and [2]School of Engineering and Materials Science, Queen Mary, University of London, Mile End Road, London, E1 4NS, UK.*

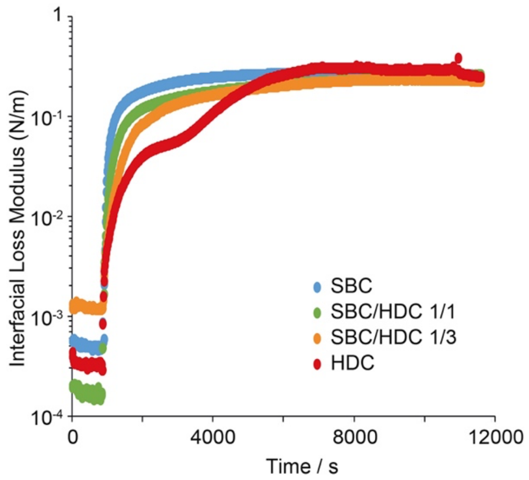


**Supplementary Figure S1. Pro-surfactant assisted PLL assembly at liquid PDMS interfaces.** Representative interfacial time sweep rheology profiles (interfacial loss moduli) recorded at a frequency of 0.1 Hz and strain of 1% ([PLL] = 100 μg/mL; [pro-surfactant] = 10 μg/mL; weight ratios are reported for mixed surfactants).

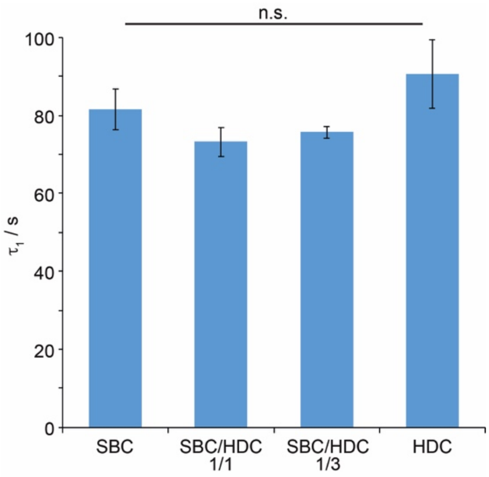


**Supplementary Figure S2. Viscoelasticity of PLL nanosheets formed at liquid PDMS interfaces.** Relaxation constants ($\tau_1$) extracted from stress relaxation profiles after double exponential fits (error bars are s.e.m.; n=3). Stress relaxation experimental details: 1% strains; [PLL] = 100 μg/mL; [pro-surfactant] = 10 μg/mL; weight ratios are reported for mixed surfactants.

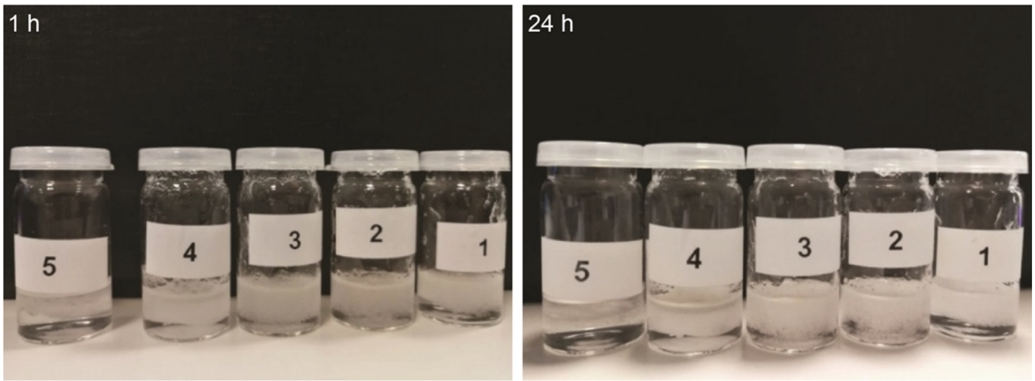


**Supplementary Figure S3. Stability of silicone oil emulsions formed with PLL nanosheets.** Pro-surfactants assisted PLL nanosheet-stabilised PDMS emulsions formed in the presence of pro-surfactant combinations (SBC/HDC at ratios of: 1, 1/0; 2, 3/1; 3, 1/1; 4, 1/3; 5, 0/1) and left for 1 h (top) and 24 h (bottom).
